# Impact of Optogenetic Activation of the Thalamic Reticular Nucleus on Sleep Architecture in Mice

**DOI:** 10.1101/2024.10.04.616683

**Authors:** Mayuko Arai, Sean Tok, Brianne A Kent

**Affiliations:** Department of Psychology, Simon Fraser University, Burnaby, Canada; Institute for Neuroscience and Neurotechnology, Simon Fraser University, Burnaby, Canada

**Keywords:** thalamic reticular nucleus, sleep, Alzheimer’s disease, optogenetics

## Abstract

Alzheimer’s disease (AD) is a progressive neurodegenerative disorder affecting millions worldwide and is often accompanied by significant sleep disturbances, such as sleep fragmentation, early awakenings, decreased sleep efficiency, and insomnia. It has been suggested that the alterations in activity of the thalamic reticular nucleus (TRN) are closely associated with sleep disruptions in AD. Evidence suggests that activating neurons expressing gamma-aminobutyric acid (GABA) within the TRN may enhance sleep quality and potentially ameliorate neuropathology associated with AD. However, the precise mechanisms through which TRN influences sleep disruptions and AD pathophysiology remain poorly understood. In this study, we investigated whether activating GABAergic TRN neurons could alter sleep architecture in wild-type mice. Utilizing optogenetic stimulation, we observed that activation of these neurons did not significantly alter sleep state durations or delta wave power, a key indicator of Slow Wave Sleep (SWS). Furthermore, the application of a two-virus strategy inadvertently led to non-specific opsin expression beyond the targeted TRN area. We discuss the potential factors that contributed to these outcomes, providing directions for future investigations to better delineate the role of the TRN in sleep and AD.

## Sleep Disruption Associated with Alzheimer’s Disease

Alzheimer’s disease (AD), a progressive neurodegenerative disorder that impacts over 58 million individuals worldwide, accounts for 60-80% of dementia cases (Alzheimer’s Association, 2024). AD is characterized by an accumulation of amyloid-beta (Aβ) containing plaques and tau-containing neurofibrillary tangles (for a review, see Breijyeh & Karaman, 2020). Sleep disturbances, prevalent in most AD patients, manifest as sleep fragmentation, early awakenings, reductions in non-rapid eye movement (NREM) and rapid eye movement (REM) sleep, and increased daytime napping (for a review, see Kent et al., 2021). Recent studies underscore the potential of sleep disturbances to exacerbate AD progression by altering the dynamics of Aβ production and clearance, along with the accumulation of abnormally phosphorylated tau protein (Barthélemy et al., 2020; Lucey et al., 2018; Winer et al., 2019). These findings are also supported by studies in mouse models of AD, which link sleep disruption to increased Aβ and phosphorylated tau levels, often considered the pathogenic drivers of AD (for a review, see Kent et al., 2021). Despite this accumulating evidence, the precise mechanisms through which sleep disruptions contribute to AD pathophysiology remain elusive. Understanding these pathways is crucial for developing targeted interventions that could mitigate the progression of AD through the treatment of sleep disturbances.

## The Role of the Thalamic Reticular Nucleus in Sleep Regulation and Alzheimer’s Disease

It has been suggested that the thalamic reticular nucleus (TRN) may play a critical role in sleep disturbances and AD. The TRN is a subcortical structure, composed of a sheet of GABAergic neurons that release gamma-aminobutyric acid (GABA) (Crabtree et al., 2018). The TRN is known to modulate sleep architecture and associated neural oscillations through robust inhibition of the thalamus during sleep periods (Lewis et al., 2015; Visocky et al., 2023). During wakefulness, the TRN engages with sensory thalamic nuclei to facilitate selective attention (Halassa & Acsády, 2016). Recent studies suggest the TRN is also actively involved in the initiation of sleep spindles (Lewis et al., 2021; Latchoumane et al., 2017; Visocky et al., 2023). Sleep spindles are a burst of rapid, rhythmic brain activity that occurs during NREM sleep, particularly N2, and is associated with memory consolidation (Leong et al., 2022). Importantly, the TRN is believed to play a crucial role in both sleep maintenance and the regulation of slow wave sleep (SWS), which is the N3 stage of NREM sleep in humans (Jagirdar et al., 2021). SWS is critical for declarative memory consolidation (Diekelmann & Born, 2010) and Aβ clearance (Xie et al., 2013), and thus, these functions of the TRN underscore its potential impact on conditions such as AD.

The TRN exhibits anatomical and functional differences between its rostral and caudal sides. The rostral subdivision primarily receives fibers from limbic thalamic nuclei, which are not involved in sensory information propagation (Visocky et al., 2023). The dorsorostral TRN, is implicated in arousal-related activities; the activity of neurons here intensifies during arousal in wakefulness and decreases in correlation with sleep-associated oscillations, such as spindles and slow waves (Crabtree, 2018). Supporting this, a recent study demonstrated that optogenetic inhibition of the rostral TRN leads to elongated sleep episodes, suggesting its significant role in arousal mechanisms (Visocky et al., 2023). In contrast, the caudal TRN interacts with thalamic nuclei responsible for integrating sensory information (Visocky et al., 2023), and the dorsocaudal TRN is suggested to be involved in sleep-related activity, such as slow waves and sleep spindles (Crabtree, 2018). A recent study inhibiting dorsocaudal TRN region using optogenetics reported that the inhibition induced fragmented sleep (Visocky et al., 2023).

Recently, there has been growing interest in the association between TRN and AD. It is suggested that the neurons in the TRN may be present and functional, but less active in a mouse model of AD, expressing human amyloid precursor protein (APP) (Jagidar et al., 2021; Hazra et al., 2016). One study utilized excitatory Designer Receptors Exclusively Activated by Designer Drugs (DREADDs) to restore the activity of TRN neurons in APP mice (Jagidar et al., 2021).

This activation not only improved sleep architecture but also decreased Aβ plaque burden, a key neuropathological feature of AD (Jagidar et al., 2021). This emerging evidence highlights the potential role of TRN as a therapeutic target for ameliorating key aspects of AD pathology.

## Modulating TRN Activity with High Spatial and Temporal Precision Using Optogenetics

In the present study, we employed optogenetics to examine the role of TRN in sleep and sleep disruptions associated with AD. Optogenetics is a technique that controls the activity of specific types of neurons using light (Boyden, 2011). This technique combines optical and genetic methods to achieve bidirectional control of neural signaling by expressing light-sensitive proteins, known as opsins, in mammalian cells (Sidor et al., 2015). Optogenetics is recognized for its ability to precisely manipulate neuronal activity with exceptional spatial and temporal accuracy, both *ex vivo* and *in vivo* (Swanson et al., 2022).

One of the pioneering studies of optogenetics was conducted in 2005 when Boyden, Deisseroth, and their colleagues demonstrated that optogenetic stimulation of mammalian neurons could control neural activity by leveraging Channelrhodopsin-2 (ChR2) (Boyden et al., 2005). ChR2, a light-gated cation channel, was originally discovered in the unicellular green alga *Chlamydomonas reinhardtii* (Boyden et al., 2005). It was found that upon photon absorption, opsins like ChR2 and halorhodopsin (NpHR) undergo a conformational change, facilitating ion transport across the plasma membrane, which results in either the depolarization or hyperpolarization of neurons (Swanson et al., 2022).

Optogenetics does have limitations due to its invasiveness, which may restrict its application in certain animal experiments and therapeutic contexts. Notably, the technique requires precise gene and light delivery to the target area and the expression of optogenetic proteins may impact cell health (Allen et al., 2015). Additionally, the heat and light from stimulation could potentially alter the physiology of local and distant circuits (Allen et al., 2015). The use of blue light in sleep studies might influence animal behaviour and neurological activities since mice, commonly used in optogenetic studies, have photoreceptors sensitive to blue light (Araragi et al., 2021).

Despite these challenges associated with optogenetics, measures like optimizing protein expression and calibrating light intensity can help overcome these issues. The strengths of optogenetics include the ability to modulate cell types specifically and precisely on a millisecond timescale (Boyden et al., 2015; Berg et al., 2020; Sidor et al., 2015). In addition, optogenetics offers substantial flexibility for experimental applications at the cellular, organ, or whole-animal level (Ferenczi et al., 2019). Various firing patterns of neurons can be produced by employing different stimulation paradigms (Mahmoudi et al., 2017). The activity of neurons can also be manipulated bidirectionally by using more than one type of opsins. For instance, it is possible to co-express ChR2, an excitatory opsin activated by blue light, with halorhodopsin, which is an inhibitory opsin activated by yellow light (Swanson et al., 2022). Furthermore, implanted devices for stimulation can be used repeatedly without the need for additional treatment (Berg et al., 2020). The precise control and flexibility of optogenetics enable innovative approaches, offering significant benefits in advancing the field of neuroscience.

## Objectives

Despite significant advancements in optogenetic techniques, there has been limited research using optogenetics to explore the potential connection between the TRN and sleep disruptions in AD. In the present study, we aimed to optogenetically activate the GABAergic neurons in the TRN to induce NREM sleep and enhance SWS in the cortical areas of wild-type mice. We targeted both rostral and caudal coordinates of the TRN with optogenetic stimulation and evaluated the effects on sleep architecture.

## Materials and Methods

### Animals

All experimental protocols were approved by the Simon Fraser University Animal Care and Use Committee (Protocol #1353P-22). We used a total of 15 male C57BL/6J mice, aged 2 to 4 months, from Charles River Laboratories (Senneville, Quebec), due to their genetic uniformity, reduced variability, robust health, and widespread availability, making them a standard and reliable model in neuroscience research. The mice were single-housed under a 12:12h light/dark cycle with *ad libitum* access to food and water. Out of the 15 animals, 5 were used to target the caudal TRN and 5 were used to target the rostral TRN, each group being injected with viruses to express the light-sensitive ion channel in the target area. The last 5 animals served as controls, receiving injections of sterile water and the same optogenetic stimulation.

### Stereotactic Adeno-Associated Virus (AAV) Infusions

We administered intracranial injections that included a mixture of two viruses. The first virus (rAAV-VGAT1-CRE-WPRE-hGH polyA) was used to express the Cre recombinase protein under a GABAergic promoter (VGAT1) in the TRN. The second virus (pAAV-EF1a-Chr2(H134R)-DIO-YFP) was used to express the light-sensitive ion channel, ChR2, in Cre-positive cells (i.e., GABAergic cells in the TRN). The mixture was prepared to inject a total of 250 nl per animal, maintaining a 3:7 ratio of Cre to ChR2 virus. Throughout the surgery, isoflurane was administered to maintain deep anesthesia. Intracortical viral injections were administered to the right posterior cortex. We tested the following coordinates to target the rostral and caudal TRN: Rostral AP: -0.8, ML: -1.2, DV: 3.5 and Caudal AP: -1.4, ML: -2.0, DV: 3.4, relative to bregma. A 1-ul Hamilton syringe was used to inject either the virus mixture or sterile water as the control. Following the injection, the virus was allowed to diffuse for 5 minutes at each coordinate, and then for an additional 5 minutes after the needle was moved up by 0.2 mm. For analgesia, Meloxicam (5 mg/kg, IP) and lidocaine (7 mg/kg, SC at the incision site) were administered at the start of the procedure. To prevent dehydration, Lactated Ringer’s solution (10 mg/kg) was given at the end of surgery. All animals were given at least 5 weeks of incubation time to allow for viral expression before initiating optogenetic stimulation.

### EEG and Optical Fiber Implantation

Following the virus incubation periods, the mice were implanted with 2-channel electroencephalogram (EEG)/ 1-channel electromyography (EMG) head caps (Pinnacle Technology, catalog number 8201-SS) and fiber optic cannulas. During the implantation surgery, the animals were maintained under deep anesthesia using isoflurane. First, a fiber optic cannula (250μm diameter, NA 0.66) from Prizmatix (Holon, Israel) was inserted into the right TRN, targeting the rostral or caudal TRN (Rostral AP: -0.8, ML: -1.2, DV: 3.5; Caudal AP: -1.4, ML: - 2.0, DV: 3.4, relative to bregma), using a custom-designed stereotaxic arm. A metallic cannula cover was attached to each optic fiber to prevent light leakage. The fiber was then secured with a small amount of vet bond (3M, London, ON) and light-curing adhesive (Pentron, Orange, CA). Once the adhesive was cured, EEG/EMG headmounts were implanted. The EEG implant involved placing four stainless steel screws at coordinates AP: +/-3 mm, ML: +/-1.5 mm relative to bregma. These screws were inserted through a prefabricated EEG headmount.

Additionally, two EMG electrode wires were placed under the nuchal muscles. Dental cement mixed with black acrylic paint (to prevent light leakage) was applied to firmly secure both the optic fiber and the EEG headmount. For analgesia, Meloxicam (5mg/kg, IP), buprenorphine (0.07mg/kg, SC), and lidocaine (7mg/kg, SC at the incision site) were administered at the start of the procedure. To prevent dehydration, Lactated Ringer’s solution (10mg/kg) was administered at the end of surgery. A recovery period of at least seven days was allowed before the commencement of EEG recordings.

### EEG Recordings and Optogenetics Stimulation

After a recovery period of one week, EEG/EMG signals were recorded for 3 hours from Zeitgeber Time (ZT) 14 as a baseline. Following this baseline assessment, the mice were optogenetically stimulated with a 473 nm LED (blue light). Three different stimulation patterns, based on previous studies (Lewis et al., 2015; Ni et al., 2016; Viscoky et al., 2023), were employed. All stimulations started at ZT14 and continued for 3 hours. For chronic stimulation, a 30-second stimulation period was alternated with a 30-second no-stimulation period over the 3 hours. The other two stimulation patterns were tonic, in which either 3 Hz or 8 Hz 10 ms pulses were repeated for 1 second, followed by a 6-second off period. The light output for each animal was adjusted to achieve a power of 4.5 mW, based on the power measurements taken for each optic fiber before surgery. To minimize light leakage from the optogenetic cannula, the entire optogenetics cable was covered with a heat shrink tube. The EEG cables were lengthened to allow simultaneous EEG recording and optogenetic stimulation. Video cameras enabled 24-hour monitoring of the animals and enhanced the accuracy of sleep scoring based on the EEG/EMG data.

### Sleep Analysis

EEG recordings were analyzed using Sirenia Sleep Pro software (Pinnacle Technology). EEG recordings were evaluated for wake, NREM, and REM sleep stages using 10-second epochs. Initially, the data were clustered by grouping epochs according to EEG and EMG frequency bands (e.g., delta, theta, alpha, beta, gamma), categorizing periods of NREM, REM, and Wake. The classification of each epoch was subsequently verified by visual review of EEG and associated video recordings, along with spectral plots. Wake periods were characterized by low-amplitude EEG (predominant frequency above 4 Hz) and high-amplitude EMG, while NREM sleep was characterised by high-amplitude EEG with frequencies under 4 Hz and low-amplitude EMG. REM sleep was identified by predominant EEG frequencies ranging between 4 and 8 Hz, consistent low-amplitude EEG waveforms, low-amplitude EMG, and a transition from NREM. Epochs were classified based on the predominant state (>50%) within each 10-second epoch.

Power spectrum analysis utilized the Fast Fourier Transform. Frequency bands were defined as follows: delta (1–4 Hz), theta (4–8 Hz), alpha (8–12 Hz), beta (12–30 Hz), and gamma (30–50 Hz), with a bandpass filter applied from 1-100 Hz to all data to remove low-frequency artifacts below 1 Hz.

### Histology

To assess viral expression, animals were perfused with phosphate-buffered saline (PBS), and brains were extracted and fixed in 4% paraformaldehyde (PFA). After 24 hours, the brains were placed in a 30% sucrose solution for 48 hours, then embedded in optimal cutting temperature (OCT) gel (Sakura Finetek USA, Torrance, CA). Coronal and sagittal slices, 40 microns thick, were collected and stored in PBS. Sections were blocked with Normal Goat Serum (Vector Laboratories, Brockville, ON), followed by incubation with Anti-VGAT Polyclonal Antibody (1:1000; catalog # PA5-27569; Thermo Fisher Scientific, Waltham, MA). Subsequently, the slices were incubated with Goat Anti-Rabbit IgG (H+L) Cross-Adsorbed Secondary Antibody, Alexa Fluor™ 647 (1:1000; catalog # A-21244; Thermo Fisher Scientific, Waltham, MA). Images were captured on a confocal microscope to display DAPI staining in blue, EYFP in green, and VGAT in red/purple.

### Statistical Analysis

Statistical analyses were performed using R (version 4.3.3). Power data were analyzed both raw and normalized as a percentage of the accumulated power from 1–100 Hz for each EEG electrode (frontal and parietal) to reduce inter-animal variability. Mixed-design ANOVA was used to assess differences in the time spent in each state across different stimulation paradigms. To compare data across different time points at 10-minute intervals, we employed a Linear Mixed Model (LMM) using the nlme package, incorporating an added quadratic effect to capture potential non-linear trends. The fixed effect in the model was the time from stimulation onset (ranging from 10 to 170), and animal ID was included as a random intercept. The largest model for the dependent variable followed the formula: DV ∼ time + time^2^. LMMs are particularly effective at accounting for both fixed effects, such as the influence of time, and random effects, such as individual differences among animals. This approach enabled us to accurately assess the impact of time on the dependent variable—deviation from baseline—while effectively managing variability introduced by individual animal differences.

## Results

### Sleep-Wake Durations During 3-Hour Stimulation

The 3 hours of EEG recordings taken during the optogenetic stimulation were analyzed to determine the time spent in each sleep or wake state (Figure 1). For NREM sleep duration, the mixed-design ANOVA indicated a significant main effect of stimulation type (comparing 30-second chronic stimulation, 3 Hz tonic stimulation, and 8 Hz tonic stimulation) (F(3, 30) = 4.00, p = 0.016). There were no significant effects for injection conditions (comparing rostral, caudal, and control) (F(2, 10) = 1.78, p = 0.218) or their interaction (F(6, 30) = 0.54, p = 0.774), suggesting no significant difference across injection sites. Post hoc analysis using Tukey HSD showed no significant pairwise differences among stimulation types (p > 0.05). Similarly, for wake duration, a significant main effect of stimulation type was observed (F(3, 30) = 3.02, p = 0.045), but no significant effects for injection conditions (F(2, 10) = 1.67, p = 0.237) or their interaction (F(6, 30) = 0.65, p = 0.690). Post hoc analysis also did not show any significant pairwise differences among stimulation types (p > 0.05). For REM sleep, no significant effects were found for stimulation type (F(3, 30) = 2.727, p = 0.062), injection coordinate (F(2, 10) = 0.226, p = 0.802), or their interaction (F(6, 30) = 1.502, p = 0.211). These findings suggest that different stimulation paradigms and injection conditions did not significantly affect the durations of sleep or wake states within the study’s constraints.

**Figure 1.**
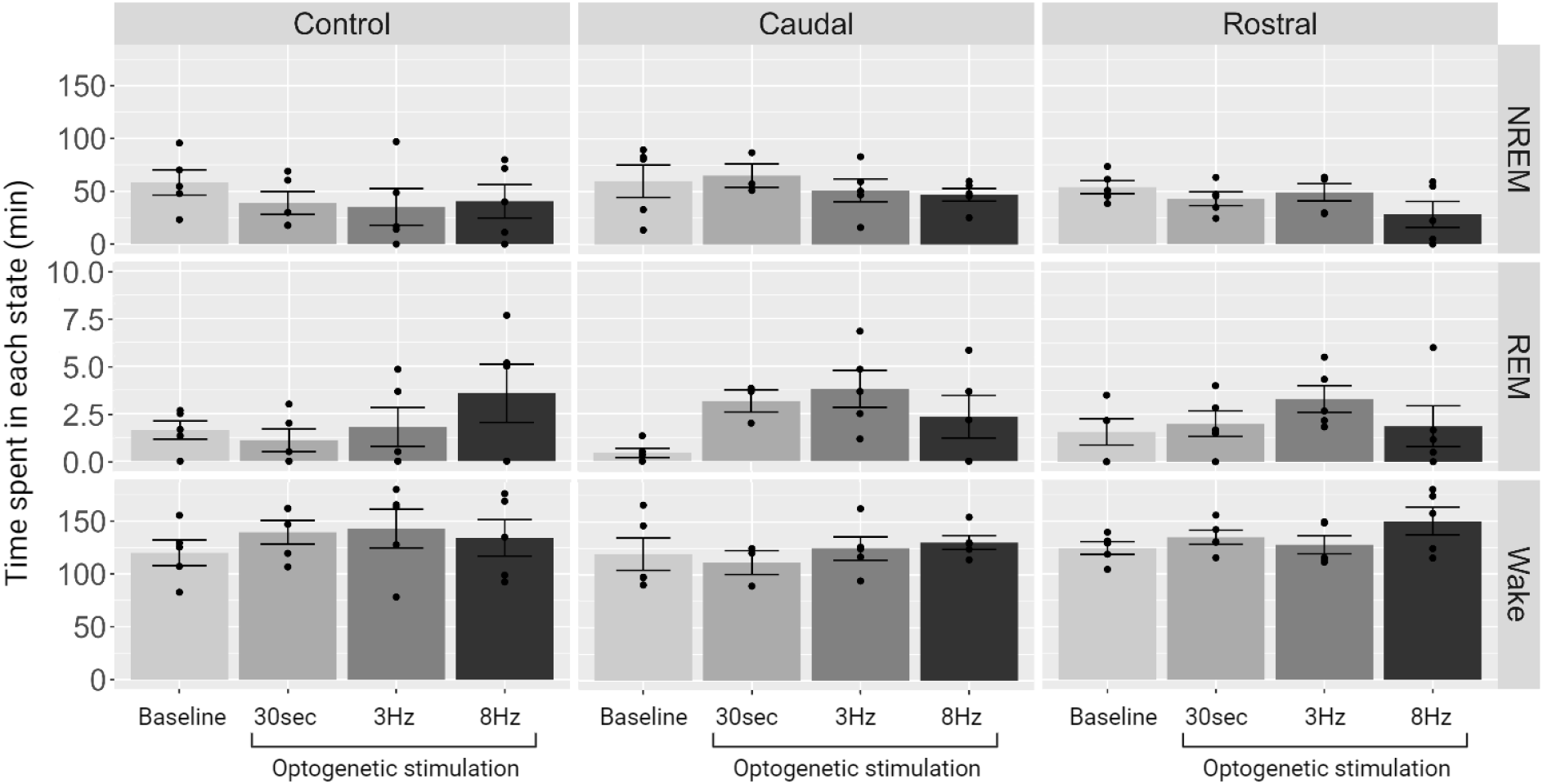
Time spent in each state during the optogenetic stimulation. The time spent in each sleep or wake state (minutes) over the three-hour optogenetic stimulation period. Graphs are presented separately for the control group and the ChR2-injected groups targeting the caudal or rostral TRN. Mean ± SEM. Black dots represent data from an individual mouse.

### Effects of 3 Hz Stimulation on NREM Duration and Delta Power at 10-Minute Intervals

To assess changes throughout the three hours of 3Hz optogenetic stimulation, the time spent in NREM sleep was analyzed every 10 minutes. When the caudal TRN was stimulated (Figure 2A), the analysis of the scaled Linear Mixed Model revealed a significant quadratic effect for the scaled time variable (coefficient = −1.08, p = 0.0150, df = 83). This negative coefficient indicates that the relationship between time and the deviation in NREM sleep duration from the baseline follows a downward curving pattern. This suggests that as time progresses from the onset, the deviation in NREM sleep duration initially increases but then decreases, forming a parabolic trend.

**Figure 2.**
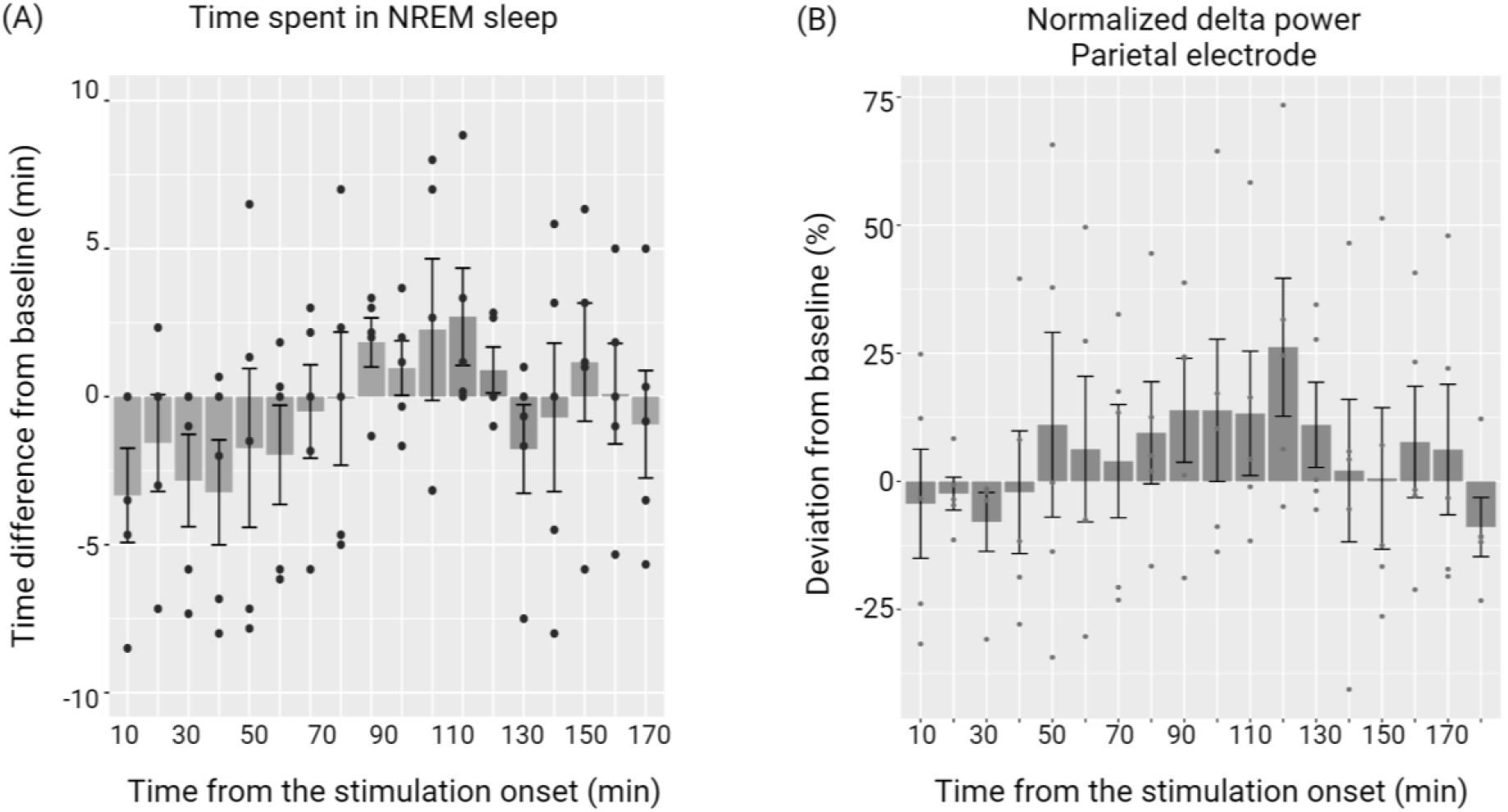
Changes in the time spent in NREM sleep and normalized delta power during 3Hz stimulation targeting the caudal TRN. (A) Differences from baseline in NREM sleep time, plotted in 10-minute intervals during the same stimulation period. (B) Normalized delta power deviations from baseline in the parietal electrode, expressed as a percentage and plotted every 10 minutes. The power was averaged for each animal over 10-minute intervals, without distinguishing between wake and sleep states. Mean ± SEM. Black dots represent data from an individual mouse.

However, the stimulation did not lead to the same effects when targeting the rostral TRN or in the control group. For the rostral TRN condition, the quadratic term had a coefficient of -0.48 (p = 0.2642, df = 83). For the control group, which did not receive the viral vector injection, the quadratic term had a coefficient of 0.12 (p = 0.7976, df = 83). These results indicate that the time-related changes in NREM duration did not exhibit a significant quadratic pattern in these conditions.

Additionally, the normalized delta power spectra were analyzed without distinguishing between sleep or wake states. The deviation from the baseline in normalized delta power, expressed as a percentage of the baseline power, was examined during the 3 Hz optogenetic stimulation. When the caudal TRN was stimulated, the scaled Linear Mixed Model for the deviation in normalized delta power from baseline in the parietal EEG electrode revealed a significant quadratic effect of the scaled time variable (coefficient = -7.28, p = 0.0078, df = 83) (Figure 2B). This suggests a non-linear relationship, where the deviation in delta power initially increases and then decreases over time. However, for the frontal EEG electrode, the quadratic term for the scaled time variable was not significant (coefficient = -4.39, p = 0.0991, df = 83), indicating no strong evidence of a non-linear relationship between time and the deviation in normalized delta power from baseline in this region.

The analysis of normalized delta power for the rostral TRN stimulation group and the control group revealed no significant effects. For the rostral TRN condition in the parietal electrode, the quadratic term had a coefficient of 0.55 (p = 0.8499, df = 83). In the frontal electrode, the quadratic term had a coefficient of -1.49 (p = 0.5397, df = 83). Similarly, in the control group, the quadratic term for the parietal electrode had a coefficient of 2.06 (p = 0.5941, df = 83), and for the frontal electrode, the coefficient was -2.47 (p = 0.4331, df = 83). These results indicate that there were no significant quadratic effects of time on normalized delta power in these conditions.

Taken together, these findings suggest that during 3 Hz optogenetic stimulation targeting the caudal TRN, NREM duration exhibits a peak over time, as indicated by a significant quadratic effect. Consistently, normalized delta power—a key characteristic of NREM sleep—peaks in the parietal electrode during stimulation, while no such effects are observed in the frontal electrode. This regional specificity highlights more pronounced responses in the parietal region, which is proximal to the stimulation site, compared to the frontal region. The lack of significant effects when the rostral TRN was stimulated suggests potential differences in the effectiveness of optogenetic stimulation on modulating NREM sleep depending on the target region.

Additionally, the absence of effects in the control group, which was injected with water instead of a viral vector, indicates that the presence of the viral vector might be crucial for the observed effects.

### Expression of ChR2 Viral Vector

Figure 3 presents representative images showing the right TRN of two C57BL/6 mice injected with the ChR2 viral vector. All animals were analyzed, but not shown here. The confocal images revealed that the virus appears to have spread to the caudate putamen. Although VGAT staining (represented in purple) was expected to show a distinct pattern within the TRN, it was observed uniformly throughout the area without specific enrichment in the TRN across all analyzed mice, and this uniform VGAT distribution potentially contributed to the off-target expression of ChR2. Additionally, enlarged ventricles were observed in some of the sections from the different animals.

**Figure 3.**
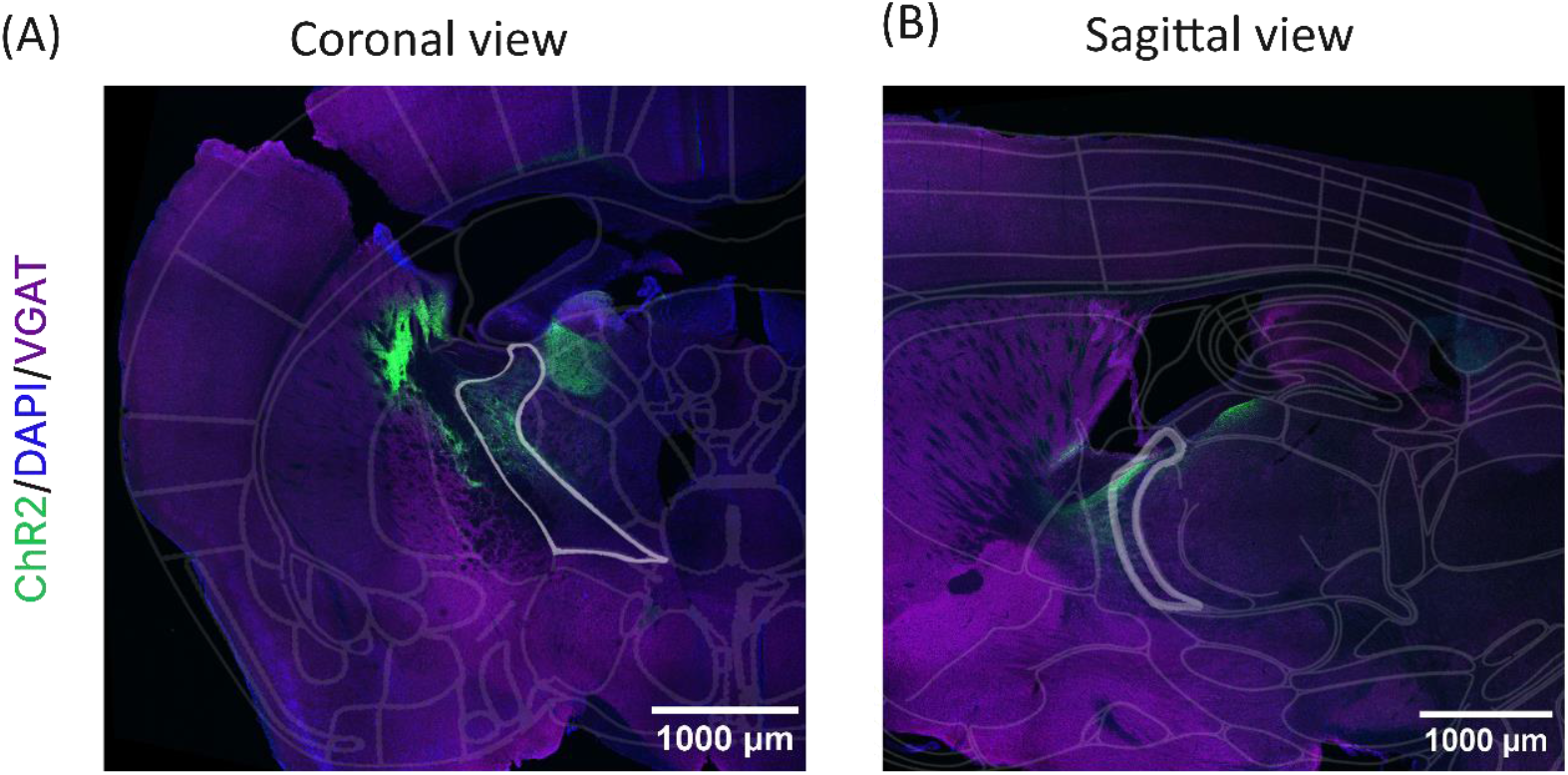
ChR2 expression in TRN. Examples of histology at 10x from C57BL/6 mice with ChR2 viral injections. The blue channel is DAPI and the green channel is EYFP, indicating off-target ChR2 expression outside of the TRN (outlined with white). Although VGAT staining in purple was expected to show a distinct pattern within the TRN, it was observed universally without specific enrichment in the TRN. (A) coronal view and (B) sagittal view.

## Discussion

Despite the potential role of the TRN in sleep disturbances associated with AD, only a few studies have investigated this possible linkage. To explore the TRN as a potential intervention target to alleviate sleep disruptions, we employed optogenetics to specifically modulate the GABAergic neurons in the TRN of C57BL/6 mice. Our results show a trend toward increased NREM duration and delta power in the parietal region during 3Hz optogenetic stimulation with an initial increase followed by a decrease, forming a parabolic pattern. However, the high variability among subjects and the off-target expression of the ChR2 viral vector warrant further investigation.

First, high variability among animals was likely due to off-target ChR2 viral expression, as illustrated in Figure 3. We used two viruses: one to deliver Cre recombinase to the target GABAergic cells, using a VGAT (vesicular GABA transporter) promoter to target these cells, and the second to deliver the opsin ChR2, which is expressed only in the presence of Cre recombinase. While the VGAT is predominantly expressed in neurons that synthesize and release GABA, the intended specific targeting of the TRN did not occur as expected. Previous studies using transgenic mice (VGAT-ChR2) observed preferential expression of ChR2 in the TRN compared to surrounding subcortical areas (Halassa et al., 2011; Lewis et al., 2015). However, in our study, VGAT expression within the TRN was not distinct and appeared to spread to the subcortical area after injecting the virus with the VGAT promoter (Figure 3). This spread of VGAT expression may have contributed to the off-target expression of ChR2 in our study. The cause of the unlocalized VGAT expression that diverges from previous findings, could be due to the specific VGAT promoter used, which varies in expression specificity. We employed VGAT1, while VGAT2 was used in the VGAT-ChR2 transgenic mice in the study by Halassa et al. (2011) (Lewis et al., 2015 did not specify). The variation in promoters could explain the off-target ChR2 expression observed, necessitating further investigation.

In addition to further investigation into the virus or transgenic mice using VGAT, future studies should also consider employing parvalbumin (PV)-Cre mice to target the GABAergic neurons in the TRN. Neurons that express PV, a Ca2+-binding protein, are believed to account for 80% of all neurons in the TRN (Vantomme et al., 2019; Visocky et al., 2023), and a subpopulation of GABAergic neurons in the TRN is suggested to express PV (Thankachan et al., 2019). Previous studies have used PV-Cre mice to selectively activate the TRN-PV neurons through viral injection to achieve Cre-dependent expression of ChR2 (Thankachan et al., 2019), or to selectively inhibit the TRN-PV neurons by achieving Cre-dependent expression of archaerhodopsin (Arch) or halorhodopsin (NpHR) (Thankachan et al., 2019; Visocky et al., 2023).

Another potential factor that may have influenced our findings relates to the limitations associated with promoting sleep in animals that do not exhibit disrupted sleep patterns. For instance, a previous study demonstrated an increase in NREM sleep duration by stimulating TRN GABAergic neurons in VGAT-Cre mice, which do not show sleep disruptions. However, the increase in NREM duration observed was modest, averaging only 3.5% (Lewis et al., 2015).

Conversely, a chemogenetic study that activated TRN GABAergic neurons using the excitatory DREADD (designer receptor exclusively activated by designer drugs) hM3Dq in APP mice—a mouse model of AD with sleep fragmentation—showed an improvement in sleep architecture. Specifically, they observed an increase in NREM/SWS duration and total sleep duration in APP mice (Jagirdar et al., 2021). Yet, in non-transgenic (NTG) mice expressing hM3Dq in the TRN, the same clozapine N-oxide (CNO) treatment did not alter SWS duration or wake time (Jagirdar et al., 2021). These findings suggest the presence of ceiling effects, or homeostatic mechanisms which are not impaired in regular, non-diseased mice. Ceiling effects may prevent significant increases in NREM/SWS, even when TRN GABAergic neurons are activated. This implies that restoring activity in the TRN could potentially improve sleep architecture in models exhibiting disrupted sleep, but not necessarily in healthy animals.

Furthermore, the power of the optogenetic LED may have also impacted our results. The enlarged ventricles observed in the histology images (Figure 3) could be attributed to the light power used in this study, as well as the reaction to the virus or inter-mouse variability. If the light power was too intense, it is possible that the surrounding area of the cannula was damaged, leading to the enlarged ventricles. However, it is also suggested that the stimulation power must reach a certain level to induce detectable changes, particularly since our 3-channel EEG/EMG system measures EEG signals only broadly in the frontal and parietal regions, making it difficult to detect changes in small local areas. Lewis et al. (2015) demonstrated that tonic optogenetic stimulation of GABAergic TRN neurons induced cortical slow waves and modulated sleep architecture. They found that when the laser power was low (<2mW), the delta (1-4 Hz) power increased only at recording sites near the ipsilateral somatosensory cortex. Conversely, when the laser power was high (>2mW), delta power increased across a broader cortical area, including the frontal and contralateral cortices. They concluded that activation of a small population of TRN neurons with weak laser power induces slow waves in a local ipsilateral cortical region, while stronger activation of a larger population induces global cortical slow waves. Their findings suggest that significant changes in EEG delta power in our study may depend on the laser power used for stimulation. It is also important to note that although Lewis et al. observed increased delta power across multiple cortical areas with high laser power (2-3.8mW), the power required to achieve similar results in the present study may have varied due to differences in ChR2 expression levels.

Lastly, it is also possible that some findings from previous studies might not necessarily indicate the beneficial effects of TRN activation on sleep due to potential issues with experimental design. For instance, Ni et al. (2016) reported that phasic spindle-like optogenetic stimulation— administered at 8Hz for 1 second at 6-second intervals for 1 hour—significantly reduced wake duration and accelerated sleep onset. They also reported that this stimulation also significantly increased the durations of total and NREM sleep during the stimulation period, compared to unstimulated controls. However, this difference could be attributed to a potential experimental design issue in Ni et al.’s study, where the control group was not optogenetically stimulated.

Ideally, the comparison should have been made with control mice that did not express optogenetic receptors but received the same light stimulation to eliminate potential confounding effects caused by the optogenetic light.

The present study underscores the need for further investigation to assess the impacts of TRN activation on sleep. Optimizing ChR2 expression using transgenic Cre-mice and refining the stimulation parameters should enable us to determine whether site-specific activation of TRN neurons indeed modulates sleep. Additionally, employing AD mouse models, which exhibit disrupted sleep, may be necessary to induce detectable changes in sleep architecture through stimulation. Further exploration of this topic will enhance our understanding of the role of the TRN in sleep and AD pathology and its potential as a therapeutic target in the future.

## Acknowledgements

We would like to thank the grant funding that made this research possible: NSERC Discovery Grant (BAK), Canadian Foundation for Innovation (BAK), Canada Research Chair (BAK), and Undergraduate Student Research Award (MA). We would also like to thank Taha Yildirim for his help with this research.

## References

Allen, B. D., Singer, A. C., & Boyden, E. S. (2015). Principles of designing interpretable optogenetic behavior experiments. Learning & Memory, 22(4), 232–238. http://www.learnmem.org/cgi/doi/10.1101/lm.038026.114

Alzheimer’s Association. (2024). 2024 Alzheimer’s Disease Facts and Figures. Alzheimer’s Association. https://www.alz.org/media/documents/alzheimers-facts-and-figures.pdf

Araragi, N., Alenina, N., & Bader, M. (2021). Carbon-mixed dental cement for fixing fiber optic ferrules prevents visually triggered locomotive enhancement in mice upon optogenetic stimulation. Heliyon, 8(1), e08692. 10.1016/j.heliyon.2021.e08692

Barthélemy, N. R., Liu, H., Lu, W., Kotzbauer, P. T., Bateman, R. J., & Lucey, B. P. (2020). Sleep deprivation affects tau phosphorylation in human cerebrospinal fluid. Annals of Neurology, 87(5), 700–709. 10.1002/ana.25702

Berg, L., Gerdey, J., & Masseck, O. A. (2020). Optogenetic manipulation of neuronal activity to modulate behavior in freely moving mice. J Vis Exp, 164. 10.3791/61023

Boyden, E. S. (2011). A history of optogenetics: The development of tools for controlling brain circuits with light. F1000 Biology Reports, 3, 11. 10.3410/B3-11

Boyden, E. S., Zhang, F., Bamberg, E., Nagel, G., & Deisseroth, K. (2005). Millisecond-timescale, genetically targeted optical control of neural activity. Nature Neuroscience, 8(9), Article 9. 10.1038/nn1525

Breijyeh, Z., & Karaman, R. (2020). Comprehensive Review on Alzheimer’s Disease: Causes and Treatment. Molecules, 25(24), Article 24. 10.3390/molecules25245789

Crabtree, J. W. (2018). Functional Diversity of Thalamic Reticular Subnetworks. Frontiers in Systems Neuroscience, 12. https://www.frontiersin.org/articles/10.3389/fnsys.2018.00041

Diekelmann, S., & Born, J. (2010). The memory function of sleep. Nature Reviews Neuroscience, 11(2), 114–126. 10.1038/nrn2762

Ferenczi, E. A., Tan, X., & Huang, C. L.-H. (2019). Principles of Optogenetic Methods and Their Application to Cardiac Experimental Systems. Frontiers in Physiology, 10. https://www.frontiersin.org/articles/10.3389/fphys.2019.01096

Halassa, M. M., & Acsády, L. (2016). Thalamic Inhibition: Diverse Sources, Diverse Scales. Trends in Neurosciences, 39(10), 680–693. 10.1016/j.tins.2016.08.001

Halassa, M. M., Siegle, J. H., Ritt, J. T., Ting, J. T., Feng, G., & Moore, C. I. (2011). Selective optical drive of thalamic reticular nucleus generates thalamic bursts and cortical spindles. Nature Neuroscience, 14(9), 1118–1120. 10.1038/nn.2880

Hazra, A., Corbett, B. F., You, J. C., Aschmies, S., Zhao, L., Li, K., … & Chin, J. (2016). Corticothalamic network dysfunction and behavioral deficits in a mouse model of Alzheimer’s disease. Neurobiology of aging, 44, 96–107. 10.1016/j.neurobiolaging.2016.04.016

Jagirdar, R., Fu, C.-H., Park, J., Corbett, B. F., Seibt, F. M., Beierlein, M., & Chin, J. (2021). Restoring activity in the thalamic reticular nucleus improves sleep architecture and reduces Aβ accumulation in mice. Science Translational Medicine, 13(618), eabh4284. 10.1126/scitranslmed.abh4284

Kent, B. A., & Mistlberger, R. E. (2017). Sleep and hippocampal neurogenesis: implications for Alzheimer’s disease. Frontiers in neuroendocrinology, 45, 35–52. 10.1016/j.yfrne.2017.02.004

Latchoumane, C.-F. V., Ngo, H.-V. V., Born, J., & Shin, H.-S. (2017). Thalamic Spindles Promote Memory Formation during Sleep through Triple Phase-Locking of Cortical, Thalamic, and Hippocampal Rhythms. Neuron, 95(2), 424-435.e6. 10.1016/j.neuron.2017.06.025

Lewis, L. D., Voigts, J., Flores, F. J., Schmitt, L. I., Wilson, M. A., Halassa, M. M., & Brown, E. N. (2015). Thalamic reticular nucleus induces fast and local modulation of arousal state. elife, 4, e08760. 10.7554/eLife.08760

Leong, C. W. Y., Leow, J. W. S., Grunstein, R. R., Naismith, S. L., Teh, J. Z., D’Rozario, A. L., & Saini, B. (2022). A systematic scoping review of the effects of central nervous system active drugs on sleep spindles and sleep-dependent memory consolidation. Sleep Medicine Reviews, 62, 101605. 10.1016/j.smrv.2022.101605

Lucey, B. P., Hicks, T. J., McLeland, J. S., Toedebusch, C. D., Boyd, J., Elbert, D. L., Patterson, B. W., Baty, J., Morris, J. C., Ovod, V., Mawuenyega, K. G., & Bateman, R. J. (2018). Effect of sleep on overnight CSF amyloid-β kinetics. Annals of Neurology, 83(1), 197–204. 10.1002/ana.25117

Mahmoudi, P., Veladi, H., & Pakdel, F. G. (2017). Optogenetics, Tools and Applications in Neurobiology. Journal of Medical Signals and Sensors, 7(2), 71–79.

Ni, K.-M., Hou, X.-J., Yang, C.-H., Dong, P., Li, Y., Zhang, Y., Jiang, P., Berg, D. K., Duan, S., & Li, X.-M. (2016). Selectively driving cholinergic fibers optically in the thalamic reticular nucleus promotes sleep. ELife, 5, e10382. 10.7554/eLife.10382

Sidor, M. M., Davidson, T. J., Tye, K. M., Warden, M. R., Diesseroth, K., & McClung, C. A. (2015). In vivo optogenetic stimulation of the rodent central nervous system. Journal of Visualized Experiments: JoVE, 95, 51483. 10.3791/51483

Swanson, J. L., Chin, P.-S., Romero, J. M., Srivastava, S., Ortiz-Guzman, J., Hunt, P. J., & Arenkiel, B. R. (2022). Advancements in the Quest to Map, Monitor, and Manipulate Neural Circuitry. Frontiers in Neural Circuits, 16. https://www.frontiersin.org/articles/10.3389/fncir.2022.886302

Thankachan, S., Katsuki, F., McKenna, J. T., Yang, C., Shukla, C., Deisseroth, K., Uygun, D. S., Strecker, R. E., Brown, R. E., McNally, J. M., & Basheer, R. (2019). Thalamic Reticular Nucleus Parvalbumin Neurons Regulate Sleep Spindles and Electrophysiological Aspects of Schizophrenia in Mice. Scientific Reports, 9, 3607. 10.1038/s41598-019-40398-9

Vantomme, G., Osorio-Forero, A., Lüthi, A., & Fernandez, L. M. J. (2019). Regulation of Local Sleep by the Thalamic Reticular Nucleus. Frontiers in Neuroscience, 13. https://www.frontiersin.org/articles/10.3389/fnins.2019.00576

Visocky, V., Morris, B. J., Dunlop, J., Brandon, N., Sakata, S., & Pratt, J. A. (2023). Site-specific inhibition of the thalamic reticular nucleus induces distinct modulations in sleep architecture. European Journal of Neuroscience. 10.1111/ejn.15908

Winer, J. R., Mander, B. A., Helfrich, R. F., Maass, A., Harrison, T. M., Baker, S. L., Knight, R. T., Jagust, W. J., & Walker, M. P. (2019). Sleep as a Potential Biomarker of Tau and β-Amyloid Burden in the Human Brain. The Journal of Neuroscience, 39(32), 6315–6324. 10.1523/JNEUROSCI.0503-19.2019

Xie, L., Kang, H., Xu, Q., Chen, M., Liao, Y., Meenakshisundaram Thiyagarajan, O’Donnell, J., Christensen, D. J., Nicholson, C., Iliff, J. J., Takano, T., & Deane, R. (2013). Sleep Drives Metabolite Clearance from the Adult Brain. Science, 342(6156), 373–377. 10.1126/science.1241224

